# Identifying Uncertainty States during Wayfinding in Indoor Environments: An EEG Classification Study

**DOI:** 10.1101/2021.12.14.453704

**Authors:** Bingzhao Zhu, Jesus G. Cruz-Garza, Mahsa Shoaran, Saleh Kalantari

## Abstract

The researchers used a machine-learning classification approach to better understand neurological features associated with periods of wayfinding uncertainty. The participants (n=30) were asked to complete wayfinding tasks of varying difficulty in a virtual reality (VR) hospital environment. Time segments when participants experienced navigational uncertainty were first identified using a combination of objective measurements (frequency of inputs into the VR controller) and behavioral annotations from two independent observers. Uncertainty time-segments during navigation were ranked on a scale from 1 (low) to 5 (high). The machine-learning model, a random forest classifier implemented using scikit-learn in Python, was used to evaluate common spatial patterns of EEG spectral power across the theta, alpha, and beta bands associated with the researcher-identified uncertainty states. The overall predictive power of the resulting model was 0.70 in terms of the area under the Receiver Operating Characteristics curve (ROC-AUC). These findings indicate that EEG data can potentially be used as a metric for identifying navigational uncertainty states, which may provide greater rigor and efficiency in studies of human responses to architectural design variables and wayfinding cues.

## 1. Introduction

Spatial navigation is an essential human skill, critical for our survival. It allows individuals to use angular and linear motion as cues to monitor their position within a space [1, 2]. This skill is particularly important in environments that are complex or novel, such as hospital buildings. In these spaces, visitors and patients often cannot build on existing experiences or expectations, and must instead rely on our spatial navigation abilities to reach a destination.

The sequence of decisions that comprise human navigation are of ten undertaken under conditions of both uncertainty and urgency, and such decisions rarely match the rational ideal for optimized path-finding. Building users may lack the spatial/cognitive abilities to interpret all of the available information about the environment with complete accuracy, and they may encounter incongruent and conflicting information that does not match other sense perceptions. Given the complexity of such facilities and the limitations of human cognition, it is unlikely that it will ever be possible to completely eliminate experiences of uncertainty and the resulting inefficient behaviors in human navigation. Thus, it is important to understand how people experience wayfinding uncertainty and how they resolve those uncertainty states.

Our understanding of exactly what happens in the brain during times of wayfinding uncertainty is currently very limited. It is well established that navigational uncertainty is usually experienced as an undesirable state, associated with discomfort and negative emotions [3, 4]. In the broader context persistent conditions of uncertainty have been linked to the emergence of sub-optimal decision strategies, as well as diminished well-being and even psychopathology [5, 6, 7, 8, 9, 10, 11].

[12] found that the type of information source (GPS device vs. human informant) influenced the decisions that participants made in situations of navigational uncertainty. The needs that people have during such conditions may differ from ordinary navigation; for example, [13] suggested that a wayfinder in uncertain conditions will eventually enter a “defensive” wayfinding mode that involves proceeding cautiously and investing excessive mental effort in scanning for conflicting information. Currently the “defensive wayfinding” model remains conceptual and largely informal, and like the overall understanding of wayfinding uncertainty it needs to be grounded in more empirical research to understand the specific neurological responses that are involved.

### 1.1. Behavioral uncertainty measurement in wayfinding studies

Clear uncertainty measurements are needed to rigorously analyze how uncertainty affects cognitive behavior [14]. Researchers have taken diverse approaches to this topic. For example, [15] used the behavioral pattern of “looking around” as an indicator of navigational uncertainty, and instrumentalized that behavior based on participant’s head motions. [16] used a more detailed “entropy value” to measure navigational uncertainty states, which the based on the purposefulness of physical motions and the extent to which participants were looking at near objects vs. far objects. [8] extensively theorized this concept of entropy, and their work has been adopted by various researchers to develop measures of uncertainty using behaviors such as walking speed, specific eye movements, and other physiological and neurophysiological responses [17, 18, 19, 20].

The predominant outlook is that wayfinding entropy arises when there is conflict between various forms of perceptual information and various behavioral options [8]. As proposed by Hirsh and colleagues, affective responses to uncertainty are linked to four primary mechanisms. First, uncertainty is a challenge that decision-makers are constantly seeking to reduce. Second, conflicts between expected outcomes and environmental cues contribute to uncertainty states. Third, expertise in a domain of endeavor can assist in resolving uncertainty. Finally, the experience of uncertainty leads to anxiety, which has measurable physiological components. This outlook provides a framework within which behaviors and measurements associated with uncertainty can be clearly defined.

Researchers have shown that uncertainty increases cognitive load, and that it often engages working memory resources, increasing vigilance and information-gathering [17, 21, 22, 23]. It also appears to promote “metacognitive” processing, in which ambiguity is overtly recognized and neural responses are activated to enhance information processing (i.e., to avoid negative consequences) [24]. Spatial navigation is likely a good domain in which to explore the cognitive impact of uncertainty more generally, given how frequently uncertainty arises during wayfinding and the importance of these processes to human survival (e.g., [15, 25]).

### 1.2. Neural dynamics of uncertainty states during wayfinding

Over the last several decades, scholars have examined the neural mechanisms associated with human spatial navigation [26, 27, 28, 29], though there has not been much particular emphasis on experiences of uncertainty in this research literature. Many of the related studies break down their findings in terms of the wayfinding strategies that are employed by participants. For example, [26] compared the use of allocentric reference frames (focused on external relationships or maps) against egocentric reference frames (focused on relationships between the environment and self) during navigational tasks and found that switching between these reference frames is mediated by the brain’s retrosplenial complex (RSC) [26, 1]. The RSC has been identified as a relevant brain region in many other studies of wayfinding, including studies on the passive viewing of navigation footage, navigations that occur mentally, and navigations in both familiar and new environments [30, 31, 32, 33, 34, 35]. The RSC is directly connected to the hippocampus as well as the occipital and parietal cortices, with indirect links to the middle prefrontal cortex [36]. These connections make it a strong candidate for being regarded as the central region for cognitive functions related to spatial orientation [37] during physical head rotations.

Functional magnetic resonance imaging (fMRI) studies have also found engagement of the parietal cortex during human wayfinding [38, 39]. In fMRI studies the activation of both the parahippocampal place area (PPA) and the RSC been seen during navigation and even during the passive observation of stimuli related to navigation [40, 41, 42, 43, 34, 35]. Many researchers believe that during spatial navigation, the PPA encodes the current environment for future recall and recognizability, while the RSC aids in orientation within the space and movements towards currently unseen navigational targets [32]. In this way, [32] asserts, the RSC and PPA have corresponding but separate roles in navigational tasks.

Another study observing the involvement of the parietal, occipital, and motor cortices in spatial navigation tasks found an association between theta-band modulation in the frontal cortex and dominant perturbations of the alpha band during navigation when participants used an egocentric reference frame. In contrast, allocentric navigation in the same study was associated with synchronization of the 12–14 Hz band and desynchronization of the 8–13 Hz band in the RSC [1]. This prior research points toward the brain regions that seem to be crucial for wayfinding and some of the EEG band dynamics that may occur during navigational tasks. However, there has been almost no research linking specific patterns that may occur in these brain regions to different wayfinding activities/sub-states such as periods of certainty vs. uncertainty.

### 1.3. Purpose of the current study

The present study was conducted to improve our understanding of neural features that may distinguish between wayfinding certainty vs. uncertainty states. We first annotated the wayfinding states (from video clips of participants in a VR hospital environment) using observational/behavioral data, and then we used a machinelearning approach to determine if those annotated states could be predicted from the participants’ EEG data. While there have been some similar recent efforts [44] in using an EEG classification approach to detect “attention states” during wayfinding, we are not aware of any other studies that have used continuous neural measures to identify wayfinding uncertainty.

We used a VR approach in this study to improve the ease of data-collection and to help reduce potential confounding variables that might impact the EEG signals and/or the ability to conduct trials (i.e., motion artifacts or potential conflicts with other individuals in the hallways) [45]. Virtual reality is a commonly used tool in wayfinding studies [46, 47, 48, 49, 50, 51, 52]. While the use of VR must be considered a limitation in terms of generalizing to real-world environments, prior research has shown that there is a strong overlap in neural responses between VR wayfinding and real-world wayfinding [53]. The use of VR also allows for a precise control of environmental design factors and precise tracking of participant behaviors [54, 55, 56], and is supported in wayfinding research [57].

The VR environment that we developed was based on actual hospital design documents. The reason for using a hospital environment in the study is that these facilities are large, complex, and unfamiliar for many visitors [58, 59]. The population that has to navigate through these complicated buildings typically includes a large number of first-time and infrequent visitors, as well as individuals who may be in a state that impairs their judgment, perception, or mobility (from sickness, anxiety, injury, etc.). Difficulties in wayfinding due to inadequate design features have been shown to be a significant source of stress for hospital patients as well as a significant burden on hospital employees and an obstacle to operational efficiency [60, 58, 61, 62]. While responses to specific architectural design features were not compared in the current study, future work using our approach may contribute to improved interior designs and more comfortable wayfinding experiences.

## 2. Materials and Methods

### 2.1. Participants

Thirty-four healthy adult participants were recruited. Data from 4 of the participants was excluded from the study due to the presence of extensive line-noise artifacts and event-logging problems. We analyzed the EEG data from the remaining 30 participants (9 reporting as female and 21 as male; M_*age*_ = 26.5, SD = 6.2, Range 20–41). After receiving verbal and written explanations of the study requirements, all participants provided written informed consent. The study procedures were approved by the Institutional Review Board for Human Participant Research (IRB) at Cornell University.

### 2.2. Procedure

The hospital environment and wayfinding tasks in this study were designed and implemented using Epic Games’ Unreal Engine. We used the Blueprints Visual Scripting system to construct the architectural environment, which was then rendered to the participants through an HTC Vive Pro head-mounted display. A non-invasive EEG cap was used to record electrical brain activity at 512 Hz for 128 channels, through the Actiview System (BioSemi Inc., Amsterdam, Netherlands) with Ag/AgCl active electrodes. The VR environments, EEG data, and experiment event marker data were timestamped, streamed, recorded, and synchronized using the Lab Streaming Layer [63].

Sessions were conducted for one participant at a time. During each session, after providing consent the participant was carefully fitted with the physiological sensors by trained research team members. To establish restingstate data, the participant was asked to sit quietly facing a blank computer monitor for one minute, and then to sit quietly with eyes closed for one minute. Once the resting-state data were collected, the participant was fitted with the VR headset and entered the virtual environment. An initial five-minute “free” period in the VR allowed the participant to become familiar with the navigational tools and to explore the platform.

During the following experiment, the same ten navigational tasks were assigned to each participant. These involved standard hospital visitor wayfinding experiences, such as locating a specific patient room (see Appendix A for a full description of the navigational tasks). To promote greater immersion, each task-series began with the presentation of a written scenario, asking the participant to imagine themselves in a moderately stressful medical situation. The total time for the whole experiment for each participant was around 120–150 minutes, including the EEG set-up, learning the VR controls, completing wayfinding tasks, and short breaks between the tasks.

### 2.3. EEG data pre-processing

The EEG data were pre-processed following [52]. The EEGLAB software package [64] was used for analysis. Raw data were imported at 512 Hz and downsampled to 128 Hz. The data were then filtered between 0.1 and 50 Hz and run through the PREP Pipeline [65], which removes 60 Hz line noise and applies a robust re-referencing method to minimize the bias introduced by referencing using noisy channels. Bad channels were removed if they presented a flatline for at least 5 seconds and if the correlation with other channels was less than 0.70 [66, 67]. Time windows that exceeded 15 standard deviations were adjusted using artifact subspace reconstruction [68], based on spherical spline interpolation from neighboring channels. The data were then re-referenced to the average of all 128 channels. Rank-adjusted Independent Component Analysis (ICA) was also used to identify artifactual components via the ICLabel toolbox [69], in order to automatically remove “Muscle” and “Eye” associated components with a threshold of 0.70. The ICs were further inspected visually by the researchers to remove artifact-laden components.

### 2.4. Identifying wayfinding uncertainty epochs

To identify periods of uncertainty during the wayfinding tasks, we first segmented the VR scenes into 5-second video clips. The video clips were parsed based on the frequency of joystick button presses, which can serve as a measure of frequent routing changes and/or reviews of the environment. The clips were then also independently labelled by two human annotators, following the protocol detailed in Appendix B. This annotation involved rating the uncertainty level in each clip on a scale from 1 (lowest) to 5 (highest), using behavioral cues such as head movements (“looking around”) and changes/reversals in direction. Overall, we obtained 1270 annotated video epochs representing participant wayfinding periods. After the annotation, we performed a two-step cleaning process to select the most representative video clips and remove ambiguous classifications. For the first step, we removed the video clips in which participants were not engaged in wayfinding activities, for example if they were standing in an elevator or were encountering technical issues (these clips were given a wayfinding uncertainty rating of “0” by the annotators to mark them for exclusion). A total of 324 video clips were excluded at this phase. In the second step, we removed video clips which failed to reflect the extreme certainty (uncertainty score = 1) or uncertainty (uncertainty score 4) states. An additional 564 clips were removed during this process, which left us with a final evaluation set of 382 video epochs that were deemed to have reliable uncertainty ratings.

### 2.5. Machine learning model

Figure 1 shows the schematic overview of how uncertainty was analyzed in relation to the EEG data. After the video clips were given a behavioral uncertainty rating by the annotators, we filtered the associated EEG recordings for those time periods to extract the theta (4–8 Hz), alpha (8–12 Hz) and beta (12–30 Hz) bands, across the entire brain. After bandpass filtering, the EEG signals were decomposed using the Common Spatial Patterns (CSP) algorithm [70]. CSP is a supervised decomposition approach, which requires “ground truth” as input. The CSP algorithm finds spatial filters that maximize the differences in variance between two classes. This algorithm identifies the informative EEG patterns that are correlated to the wayfinding uncertainty states and we chose it for use in our analysis because CSP can effectively separate signal from noise. These input conditions were based on the researcher’s classification of uncertainty states during the wayfinding tasks. The CSP transformation steps were implemented using the MEG+EEG Analysis and Visualization (MNE) tools implemented in Python [71]. After CSP transformation, we selected the top 20 CSPs from each frequency band based on the absolute deviation of their eigenvalues from 0.5. We used the average power of the CSP patterns to represent the neural activity and applied a log transform to standardize the band features.

**Figure 1:**
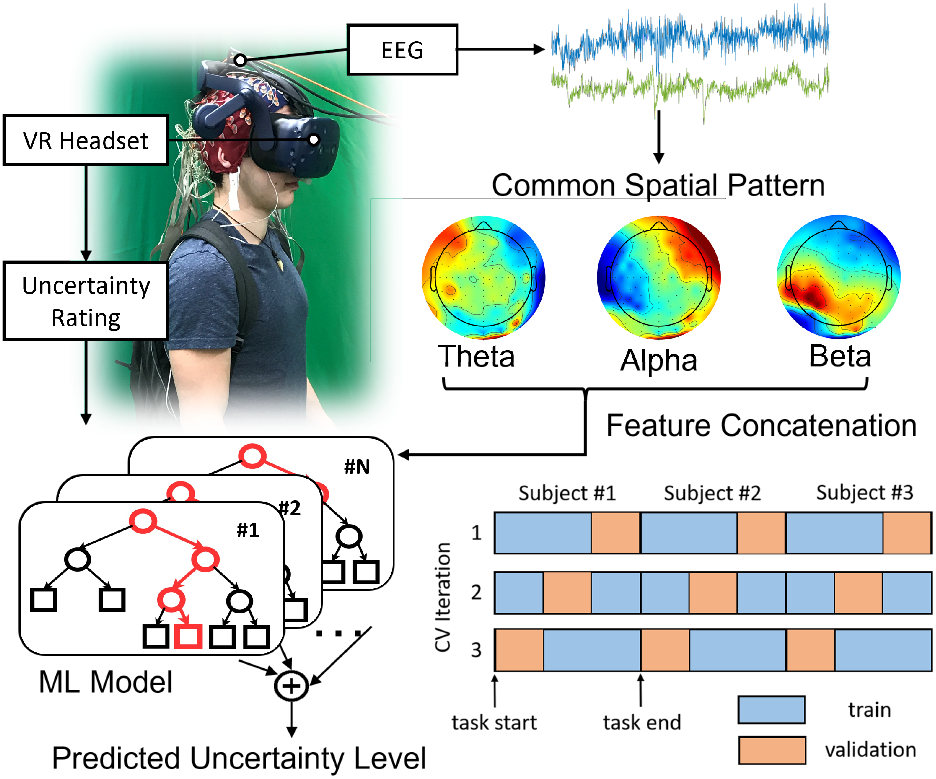
Schematic overview of decoding uncertainty during the wayfinding tasks. The certainty and uncertainty periods were first annotated by the researchers based on a rigorous screening process, described in detail in section 2.4 and Appendix B. We then extracted the common spatial pattern features from three EEG frequency bands (theta, alpha, and beta) for the annotated time epochs, and used a Random Forest classifier to identify EEG features associated with the certainty vs. uncertainty states. To evaluate classification performance, we split the EEG recordings of each subject into k-folds without shuffling. For each cross-validation (CV) iteration, we used one fold from each subject as the validation set and the other folds as the training set. This data-splitting approach is less sensitive to cross-subject differences since the training set consists of multi-subject recordings.

The features from the three bands were concatenated to construct a feature vector, which was fed into a machine learning model for classification purposes. Our goal is to predict the uncertainty state for each EEG epoch by only looking at the corresponding feature vector. We then trained a random Forest Classifier algorithm with 100 trees to predict the human-annotated uncertainty level. The classification model was implemented using scikit-learn in Python. Given the imbalanced class distribution, we measured the model’s performance in terms of the area under the Receiver Operating Characteristics curve (ROC).

To develop the machine-learning model, we separated the training and validation sets using a crossvalidation scheme as detailed in Figure 1. The EEG signals of each participant were uniformly split into five k-folds, following the chronological order of the time series. In each cross-validation iteration, we used 4 of the folds from each participant to train the model, and 1 fold from each participant for validation. As a result, both the training and validation sets included recordings from the entire group of participants, which greatly reduces the impact of cross-subject differences. The validation set consisted of a continuous, unshuffled EEG block from each subject to maintain the chronological order and minimize information leakage caused by shuffling data [72, 73, 74].

To further identify important features in the classification of uncertainty vs. certainty states, we measured the total impurity reduction contributed by each CSP. The impurity reduction is the criterion to grow decision trees and it can be efficiently calculated to quantify the importance of features in a Random Forest classifier (ensemble of decision tree). Starting from using the single attribute that achieved the highest feature importance score, we sequentially added new attributes to the subset based on their importance. With this selection approach, we were able to remove redundant CSPs and find the optimal subset to detect the human uncertainty state during the wayfinding tasks.

## 3. Results

### 3.1. Observational annotations of wayfinding uncertainty

As shown in Figure 2(a), there was a high consistency between two annotators, with a Pearson’s correlation coefficient of 0.77. Figure 2(b) shows the relation between human-annotated uncertainty scores and BPR. We observed that high uncertainty scores are associated with low BPR, where participants struggled to find the right direction and make movements. On the other hand, high BPR is associated with decisive movement which indicates low uncertainty score.

**Figure 2:**
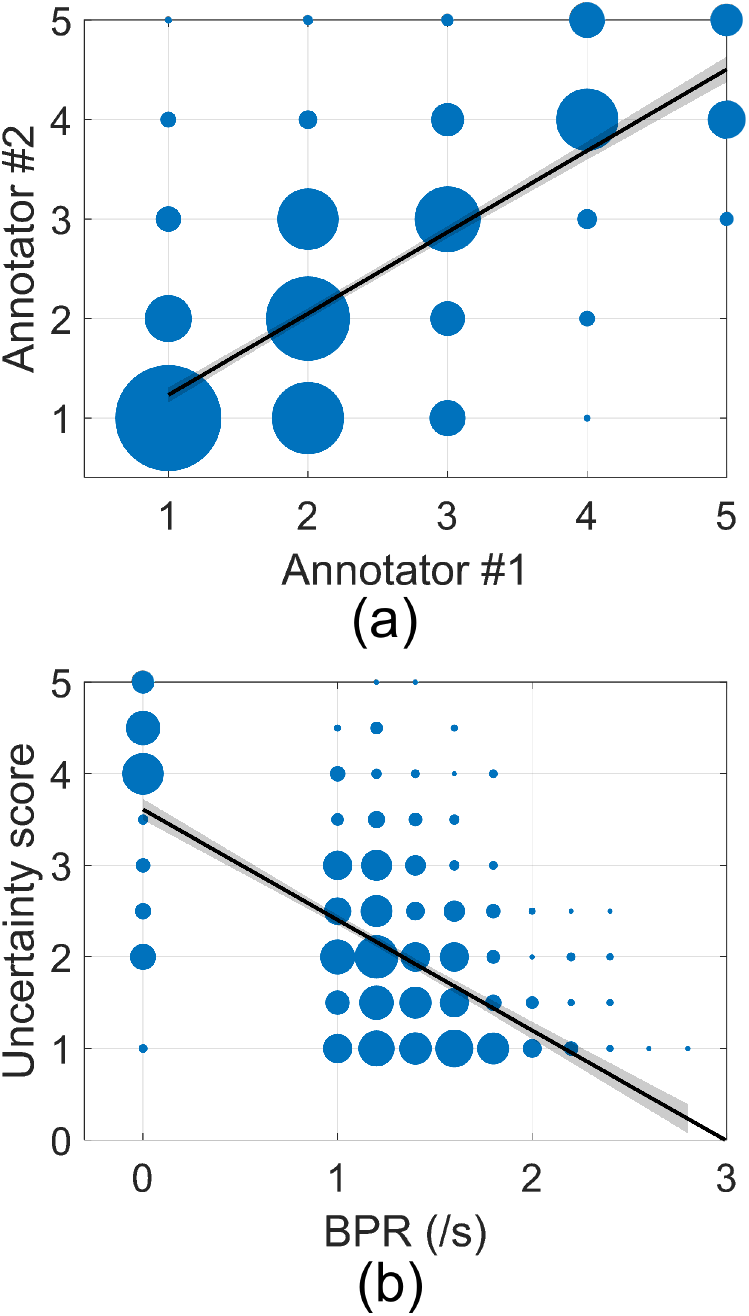
(a) Consistency between two annotators. There is a good consistency between the ratings from two annotators (Pearson’s P value: 0.77). (b) Correlation between human-annotated uncertainty scores and BPR. We observed a negative correlation between BPR and human-annotated uncertainty score (Pearson’s P value: −0.64). We show the least-squares fitted linear line with the shaded area indicating 95% confidence bound. Marker size represents the number of epochs.

### 3.2. Classification performance

Figure 3 shows the ROC for each cross-validation fold, where the mean ROC and standard deviation are indicated for the all trials. We achieved an average areaunder-the-curve score of 0.70. The classification performance is higher than the chance level (0.5), which indicates the successful distinction between certainty and uncertainty states during the hospital wayfinding task. Since we extracted all the features from EEG, our results shows that the participants’ uncertainty states can be decoded from noninvasive brain recordings.

**Figure 3:**
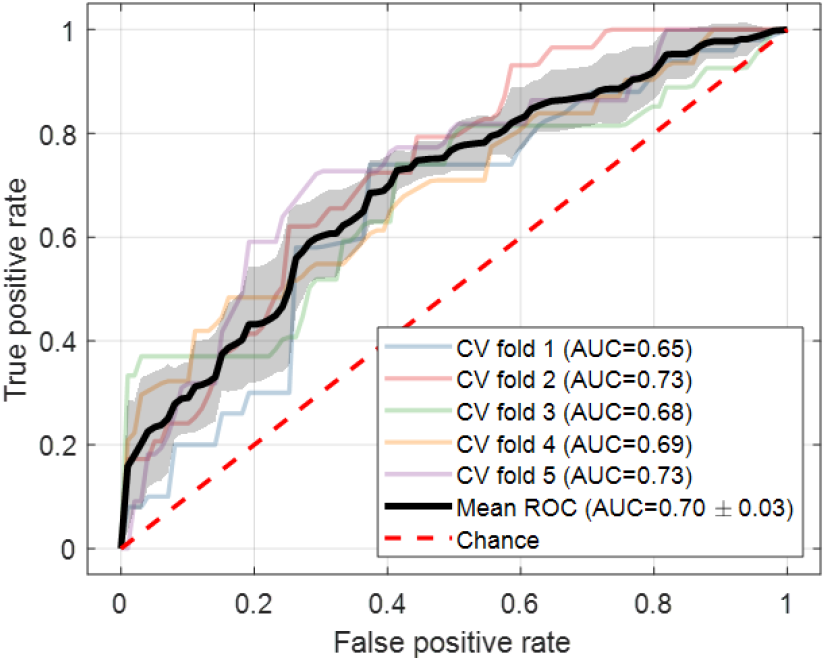
Receiver operating characteristic (ROC) curves for predicting human-annotated uncertainty scores. We show the ROC curves over 5-fold cross-validation (CV) and achieved an average area-underthe-curve score of 0.70 for uncertainty decoding. The shaded area indicates the standard deviation.

### 3.3. Feature visualization through CSPs

To better understand the informative indicators for uncertainty decoding, we interpret the model prediction using Shapley Additive Explanations (SHAP, [75]). Figure 4(a) presents 4 consecutive screen shots of a video clip, where the participant swung head and showed little intention to make a movement. This video clip received an average uncertainty score of 4.5 from the raters (i.e., very high uncertainty). With SHAP, we visualize the dominating factors which contribute to the model prediction.

**Figure 4:**
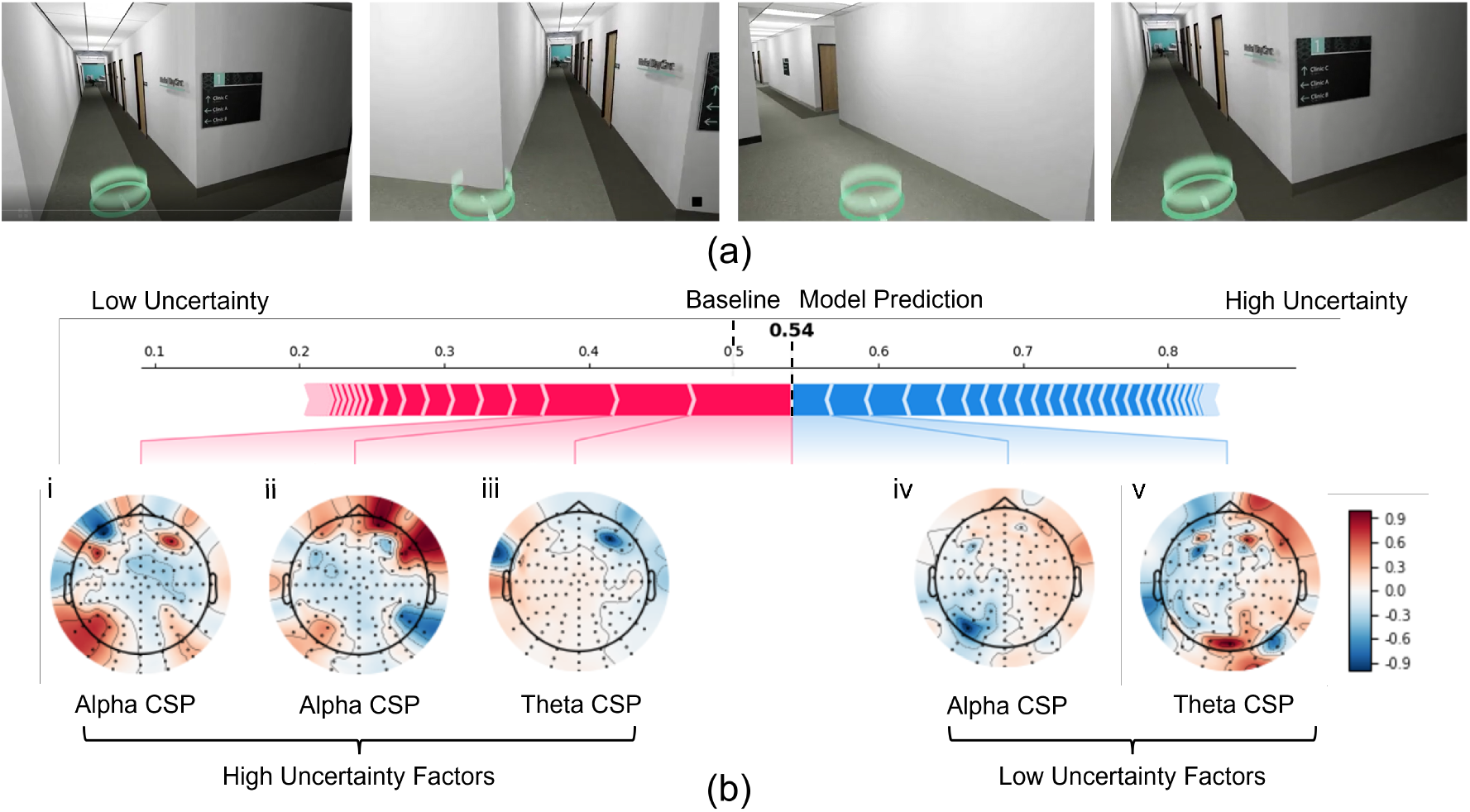
(a) Screenshots from a video clip in which the participant swung around with little intention for movement. The epoch received a 4.5 (very high) average uncertainty score from the raters. (b) Model prediction and interpretation of EEG data using Shapley Additive Explanations. The Random Forest model successfully classified the epoch as “uncertainty” by predicting an uncertainty score (0.54) higher than the baseline (0.5). The model prediction is driven by various CSP patterns from different bands. The factors contributing to high uncertainty prediction are shown in red, whereas those contributing to low uncertainty are in blue (red factors push the model prediction to the right, indicating higher uncertainty, while blue factors push the model prediction to the left). The CSP brain plots indicate that theta and alpha bands contributed most significantly to the classification of this epoch.

In Figure 4(b), red CSPs push the model to reach a high uncertainty prediction, whereas blue CSPs contributes more to a low uncertainty prediction. Overall, the red CSPs have a higher impact (indicated by the length of red/blue bars), which leads to a correct prediction of the current epoch as as a moment of uncertainty. Figure 5 is similar to Figure 4 but presents an epoch of decisive movement, which received the lowest uncertainty score (1) from both annotators.

**Figure 5:**
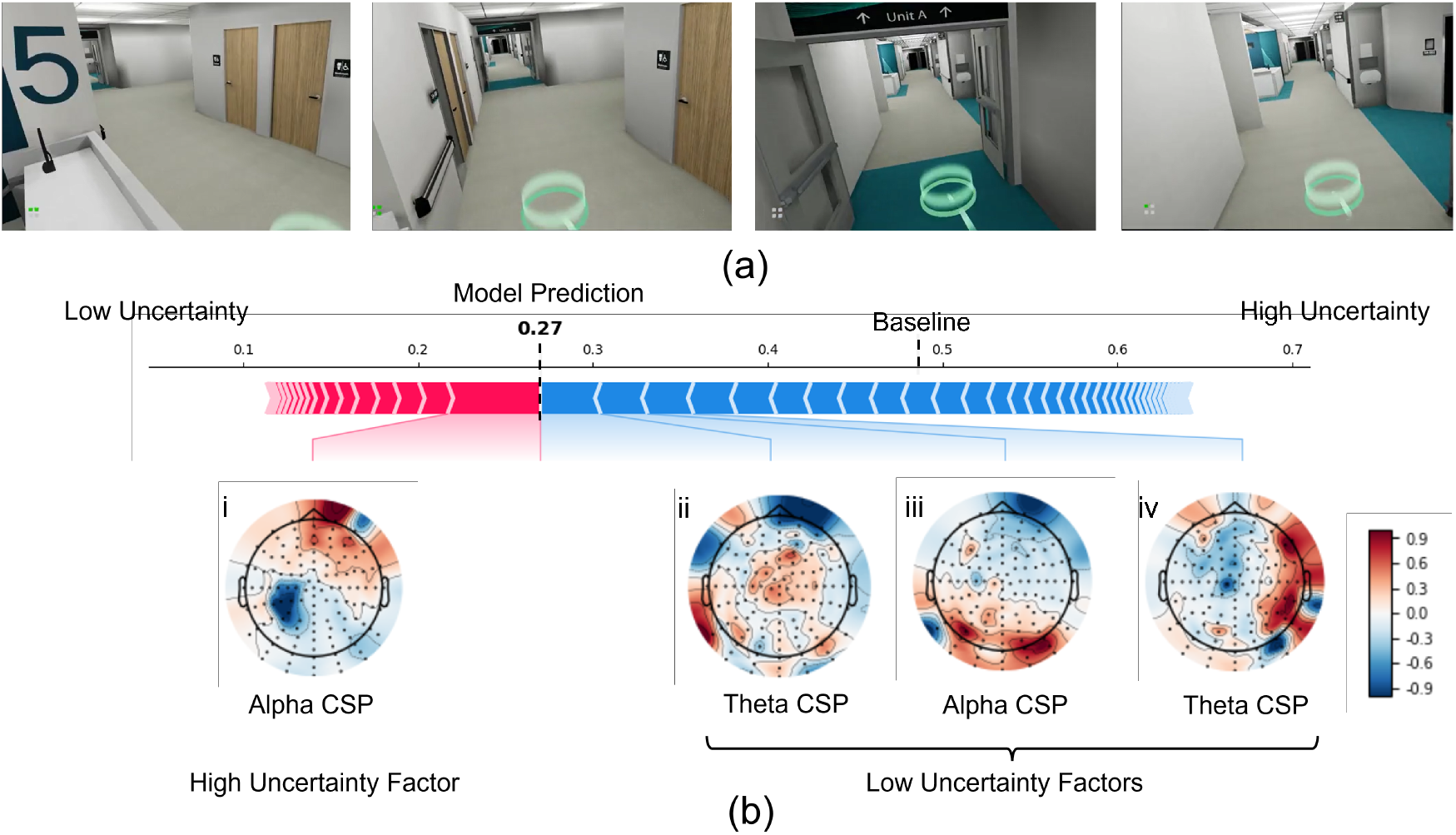
(a) Screenshot of a video clip in which the participant performed decisive movement. The uncertainty score from the raters for this clip was 1 (indicating very low levels of wayfinding uncertainty). (b) Model prediction and interpretation of the EEG data using Shapley Additive Explanations. The Random Forest model successfully classified this epoch as “certainty” by predicting an uncertainty level (0.27) lower than the baseline (0.5). The model prediction is driven by various CSP patterns from different bands. The factors contributing to high uncertainty prediction are shown in red, whereas those contributing to low uncertainty are in blue (red factors push the model prediction to the right, indicating higher uncertainty, while blue factors push the model prediction to the left). The CSP brain plots indicate that theta and alpha bands contributed most significantly to the classification of this epoch.

In Figure 4(b), an increase in alpha power in parietal-left (Fig.4(b) i) regions and frontal-right (Fig.4(b) ii) regions is associated with high uncertainty scores during wayfinding. A CSP pattern with lower theta band-power in right-frontal region (Fig.4(b) iii) contributes to the classification model prediction for the high-uncertainty class (red line). The opposite patterns were observed for the low-certainty class (blue line): there was alpha power suppression in left-parietal regions (Fig.4(b) iv), and high theta power in occipital and right-frontal regions (Fig.4(b) v).

In the example of Figure 4b, the EEG patterns associated with this epoch were classified as a “high uncertainty” because of the largest weights (red-blue line) in those patterns with increased alpha power in left-parietal and right-frontal regions (Fig.4(b) i and ii), with concurrent decreased theta power in frontal regions (Fig.4(b) iii).

Figure 5(b) shows an example of a low uncertainty navigation epoch, as classified by the EEG CSP bandpower features random forest model. This sample shows a pronounced theta power decrease in frontal and pre-frontal regions (Fig.5(b) ii), predictive of low navigation uncertainty (blue line). The most significant CSP component in alpha power for low-uncertaintly (blue line) in this example shows low weighting in frontal regions, and high weighting for occipital areas (Fig.5(b) iii), which stands in contrast to the largest contributing alpha power CSP in Figure 4(b) (ii).

Higher theta power in parietal areas has been observed in salient landmark-based wayfinding scenarios in virtual reality [50]. Increased theta power in the retrosplineal cortex has also been found when participants rotate their head searching for navigational cues in VR environments, compared to translational movement [37]. Studies in VR maze learning have also found that there are more prevalent theta episodes when a maze becomes more difficult; suggesting that increased theta activity is indicative of general demands of the task, but not necessarily associated with immediate cognitive demands [76]. In addition. Theta power increase has been positively correlated to increased task difficulty in frontal regions [1, 77, 78].

Alpha power suppression has been observed when participants maintained orientation in active translational navigation tasks [37]. During tunnel turns in VR, alpha suppression was found in visual cortex areas for egocentric-reference frame participants; while this suppression was stronger for egocentric reference-frame participants, also found in inferior parietal and retrosplineal areas [26]. Desynchronization in parietal-region alpha band appears most prominent before stimulus turns [79], while alpha power increases in right parietal areas during maintained spatial navigation [80]. Alpha suppression is associated with increased visual processing and attentional processing [81] during mobile active navigation.

### 3.4. Intrepretation of classification

Our results indicate that a small subset of CSP features can achieve a reasonably high performance in identifying wayfinding uncertainty states. Figure 6(a) shows the feature selection process, where we included the most relevant CSPs from each EEG band. The classification performance was plotted as a function of the CSP count in the subset, with the pie plots showing which frequency band the CSPs were extracted from. Using only 7 CSPs, we achieved a mean ROC-AUC score of 0.69 for predicting the wayfinding uncertainty level in each video clip, which is only 0.01 lower than was achieved by using the entire set of CSP features. Even more interestingly, these top 7 features only consist of CSPs from the theta and alpha bands, indicating that the beta band may have a limited role in the characterization of human wayfinding uncertainty states.

**Figure 6:**
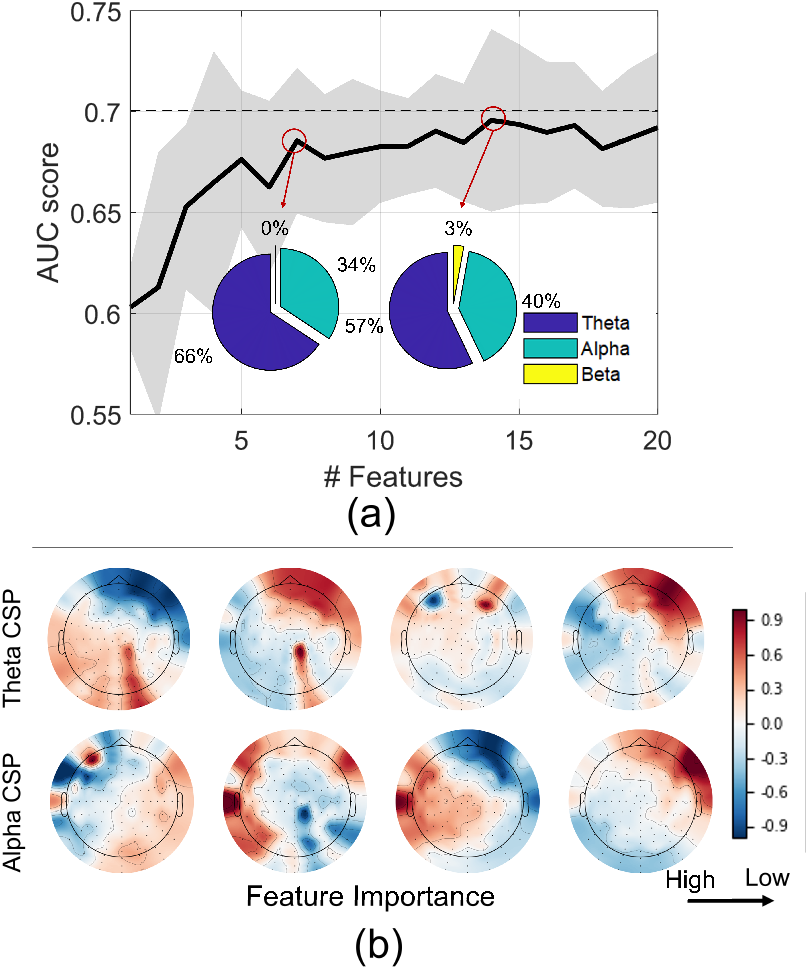
(a) Feature selection for predicting human-annotated uncertainty scores. The plot shows the classification performance as a function of the number of selected features. The baseline performance (dashed line) is achieved by using all 20 CSP features. Starting from only using one feature, we incrementally added features into the subset based on their importance. We were able to achieve the uncertainty classification with only a small subset of CSP features, with marginal performance loss. The distribution of feature types is shown by the pie plots, indicating that the most informative CSP features come from the theta and alpha bands. (b) Visualization of the most discriminative CSP patterns.

We further visualized the exact CSP patterns that are important for the wayfinding classification task. Different from the SHAP analysis which provides explanation for each epoch, Figure 6(b) visualizes feature importance from the group level. Specifically, we are interested in the CSP patterns that lead to high classification performance. Figure 6(b) shows the CSP patterns that separate the human-annotated uncertainty score extremes (i.e., uncertainty score of ≥4 vs. uncertainty score of 1) for the 5-s video clips. We observe again that the theta band and the frontal channels have most distinct variance between the certainty vs. uncertainty classes. In the alpha band the frontal and parietooccipital locations had the most significant variation, as observed with extremes in CSP weighting in these regions. In the theta band, patterns in frontal and parietal locations were also observed. These group-level weight distributions for the most discriminant CSP patterns between 5-s epochs of time where a participant navigated through the hospital setting capture the aggregate differences between uncertain and certain navigation. While Figures 4 and 5 inspect a representative epoch sample of the associated classification.

## 4. Discussion

The main goal of this study was to assess if brain activity could be used to characterize uncertainty events during navigation in a complex building environment. The results demonstrate that behavioral uncertainty in human wayfinding likely has neurophysiological correlates, which can potentially allow for the automatic classification of such uncertainty events during wayfinding tasks.

The neurophysiological interpretation of CSP patterns is only indicative of the most distinct patterns that differenciate between the annotated classes in the experiment: certain and uncertain-labeled 5-s epochs of navigation through a VR hospital setting. It is not a source localization method. The CSP algorithm finds spatial filters that maximize variance for one class while minimizing the variance for the other class [82]. A strong predictive contribution from a location in the scalp can be due to consistent potentials associated with wayfinding uncertainty, or alternatively, from consistent potentials during high-certainty navigation epochs. The patterns may also arise as a combination of both effects. CSP pattern selection at the group level (Figure 6(b)) is sensitive to outliers, as the selection is driven by eigenvalues (variance in one condition divided by the sum of variances in both conditions). The scalp map pattern visualization is constrained by these limitations. We calculated the average power of the filtered signal within each trial [82], and visualized the CSP patterns in Figures 4 and 5. These examples provide a snapshot of the random forest classifier’s decision which CSP patterns were most significant, and the discerning patterns associated with high uncertainty or low uncertainty.

In the SHAP analyses (Figures 4 and 5) there is a clear network of frontal channels in the theta band, and frontal with parieto-occipital contributions in the alpha and theta bands, that are the primary drivers of the binary classification performance. The theta band contributions in the frontal cortex mirrors previous findings of theta and alpha band involvement in active navigation. Frontal midline theta-band has been associated with active navigation in VR contexts [83], with desynchronization when an obstruction appeared. Higher theta power in parietal areas has been previously observed in landmark-based wayfinding scenarios [50] when participants evaluated the landmarks in an active navigation context. Further, in active navigational tasks, navigation based on egocentric reference frames recruited a network of parietal, motor, and occipital cortices in the alpha band, with frontal theta band modulation [1]; and retrosplineal cortex involvement in heading computation [37], but not in translational movement. Studies in VR maze learning have found that there is more prevalent theta activity when a maze becomes more difficult; suggesting that increased theta activity is indicative of general demands of the wayfinding task [76].

The current research provided the first steps in developing a continuous EEG-based measurement of wayfinding uncertainty in indoor environments. Once these neural measurements of uncertainty states are further refined and confirmed in broader studies, they can be used to conduct rigorous and efficient research with important applications for building design and pre-occupancy evaluation. The current study contributes to the development of a novel continuous measures for assessing the level of uncertainty during navigation at any given moment. As suggested by [14], continuous navigation data can provide important insights into what information someone seeks to reduce that uncertainty and can better explain the cognition-action loop contributing to spatial learning and decision making. The EEG-based classification approach to identifying wayfinding uncertainty that we developed here can potentially allows researchers to test hypotheses about the impact of environmental features on human behavior. Applications of this approach stretch across numerous architectural specialties, as well as other “spatial professions” such as the design of immersive video games and spherical cinema [84]. Continuing to improve our understanding of the neurological components of wayfinding uncertainty could also potentially contribute to new types of navigational aid design and more effective approaches to familiarizing people to a new spatial environment. In high-stakes situations, such as those involving emergency first responders or helping patients to reach the appropriate care centers, providing the right information as uncertainty arises could improve outcomes and help to reduce anxiety.

### 4.1. Limitations and Future Work

The binary classification approach followed in this study is dependent on the class labels (certainty vs. uncertainty) and the labeling procedure that was implemented. The certainty/uncertainty scores provided by human annotators followed a specific procedure (Appendix B), which may not be generalizable to other wayfinding contexts. The interpretation of the neural features a associated with the classification performance must be understood in the context of this specific rating approach, as well as the hospital environment and the types of navigational tasks performed (Appendix A).

Using VR to investigate wayfinding navigation has some limitations, particularly in that physical cues, textures, and sounds may differ from real-world environments. Some researchers have argued that the brain’s predictive capability effectively short-circuits the body and its broader related processes in VR if the visual perception is in line with the body’s actions, for instance, when head movements result in predictable alterations in visual information [85]. However, additional studies using mobile EEG in non-virtual contexts are needed to determine if the results from VR can be fully generalized to real-world environments.

Experiences of wayfinding uncertainty, along with the associated behaviors and neural dynamics, are expected to change gradually and continuously during the wayfinding process. If changes in sensoryinformation processing, decision making, and action (walking, turning, stopping) occur intermittently in a typical wayfinding task, we can expect that the associated neural dynamics would be modulated correspondingly. Our results using two-class models provide evidence of distinguishable neural features in pre-labeled certainty and uncertainty epochs, but not their modulation in transition states. In future studies we plan to conduct singletrial dynamic characterizations of behavioral and neural data, which will help to quantify the neural pattern modulations associated specific aspects of wayfinding activities and their transitions.

These effects should be studied further in regards to design elements to guide wayfinding cues in the built environment and VR spaces. Cross-participant differences and optimized machine learning models that take into account different wayfinding strategies (e.g. allocentric vs. egocentric oriented participants) [26] may provide more information about the EEG features that are linked to wayfinding certainty and uncertainty states and help to ensure that architectural designs and cues are useful for the entire human population.

Recent study [86] has shown the potential of “augmented reality” (virtual information overlayed onto real spaces) as a tool to improve wayfinding performance and decrease cognitive loads during wayfinding tasks. Findings from neurological studies on wayfinding uncertainty and responses to environmental cues may assist in the development of such tools, leading to a more context-aware and user-aware intelligent wayfinding aid system.

## 5. Conclusion

This study took a machine-learning classification approach to gain a better understanding of neurological features associated with periods of uncertainty during navigation. This study used a VR hospital environment, and participants were asked to complete wayfinding tasks of varying difficulty. Two observers independently annotated human mental uncertainty state on a scale from 1 (low) to 5 (high). We implemented random forest classifiers to predict researcher-identified uncertainty states from the EEG common spatial patterns across various frequency bands and an AUC score of 0.70. We also observed an increase in alpha power in fronto-parietal regions with a corresponding suppression of frontal theta power in high-uncertainty conditions, and the opposite patterns in the low-uncertainty condition. Our results indicate that the frontal theta and occipital alpha power of EEG can potentially be used as a metric to quantify uncertainty states during wayfinding.

## Acknowledgement

The authors gratefully acknowledge Parkin Architects, the Government of Newfoundland and Labrador, the Western Regional Health Authority and the Corner Brook Acute Care Hospital. The authors thank the team at the Design and Augmented Intelligence Lab at Cornell University, including Armin Mostafavi, Qi Yang, Talia Fishman, Viraj Govani, Julia Kan, Jeffrey Neo, Mi Rae Kim, Matthew Canabarro, Emme Wong, Clair Choi, Elita Gao, and Michael Darfler for assisting in visualization of VR environments, data collection, and labeling wayfinding uncertainty epoch. The authors also thank the interior design and wayfinding/signage design team at Parkin Architects in Joint Venture with B+H Architects.

## Appendix A. Description of Wayfinding Task

Description of the wayfinding tasks are included in the Table 2, and examples of the virtual stimuli included in the Figure A.7.

## Appendix B. Coding Wayfinding Uncertainty

We screened the recorded first-person perspective videos for all participants. Each recorded video was divided into 5-second clips, leading to a total of 1270 video segments. Wayfinding uncertainty scores were assigned to each 5-second clip using the following procedure.

First, 254 video clips were randomly selected, and two research assistants were asked to rate the navigational uncertainty of the participant during each 5-second clip, on a scale from 1 (low uncertainty) to 5 (high uncertainty), based on their own individual interpretations of the videos. The Cohen’s kappa inter-rater reliability score for these ratings was 0.48.

After this initial pilot rating, a group meeting was held to review points of consistency and divergence in the research assistants’ ratings. In this discussion we identified behavioral indicators to help the raters reduce their points of disagreement. Those behavioral indicators were: (1) decisive movement, (2) exploratory movement, (3) turning around, (4) swinging head, (5) made decision, (7) intention to move, and (8) other actions. The raters were asked to determine which of these indicators was present in each video segment.

In “decisive movement,” the participant moved without hesitation and in a firm rhythm. In contrast, “exploratory movement” referred to segments in which the participant was moving but paused frequently to evaluate signs or environmental cues to guide their navigation. Participants were “turning around” during a segment if they rotated in only one direction, from left to right for example. They were regarded as “swinging head” if they turned their heads in both directions, which implied they were hesitating. If they moved after “turning around” or “swinging head,” they were regarded as having “made [a] decision.” If they began to enter motion instructions in the controller during the video segment, then they showed “intention to move.” If the participants’ behavior during the clip was not relevant to wayfinding activities—for example if they were standing in an elevator or encountering technical issues—then the video clip would be identified as “other actions.”

The manner in which the raters were instructed to evaluate the videos is shown in Figure 9. An uncertainty score of 0 was given if the clip showed only “other actions”; these clips were excluded from the data analysis. An uncertainty score of 1 was given to videos when participants were moving decisively most of the time. A score of 2 was given if the participant was conducting exploratory movement. If the participant made a decision after turning around the video would be given a score of 3. If the participant turned around without making a decision, or if they swung their head and then initiated a movement, the video would be given a score of 4. Finally, if the participant swung their head but showed no intention to move, the clip was given an uncertainty rating of 5.

Finally, all 1270 video clips were reviewed by the two annotators. The 5-level uncertainty measurement refers to the raw ratings from annotators. We further calculated the 2-level uncertainty scores by simple thresholding (Table B.1), which is used for binary classification. The results of these final ratings produced a 0.53 kappa score for 5-level uncertainty, and 0.88 kappa score for 2-level uncertainty (Table 1).

**Figure A.7:**
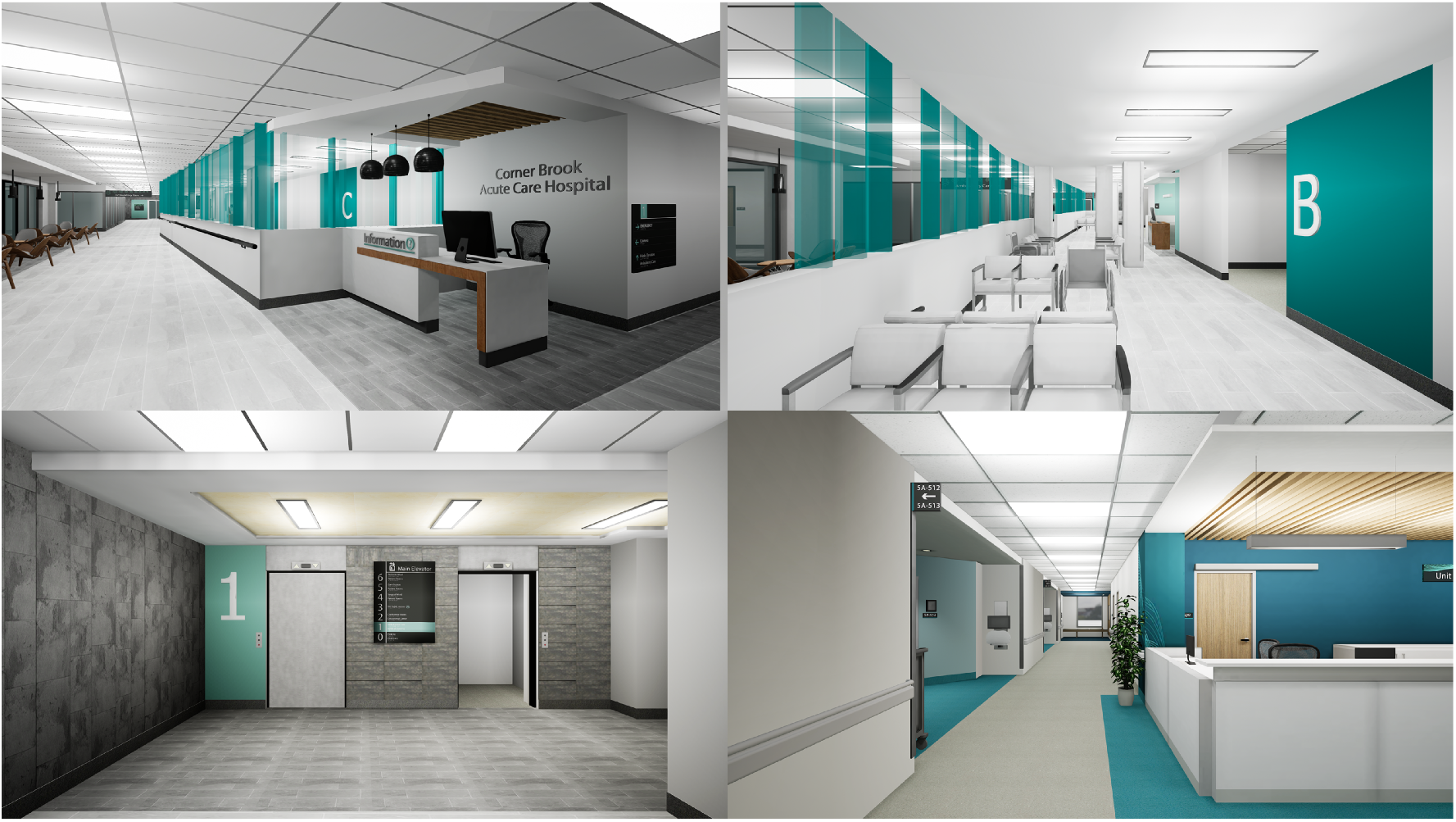
Examples (screenshots) of the VR hospital environment.

**Table B.1:** Cohen’s Kappa for the behavioral uncertainty ratings.

|  | 5-level<br>uncertainty | 2-level<br>uncertainty* | Other<br>actions |
| --- | --- | --- | --- |
| Cohen’s<br>Kappa | 0.53 | 0.88 | 0.75 |

\* uncertainty score = 1, uncertainty score $\geq 4$ .

**Figure B.8.**
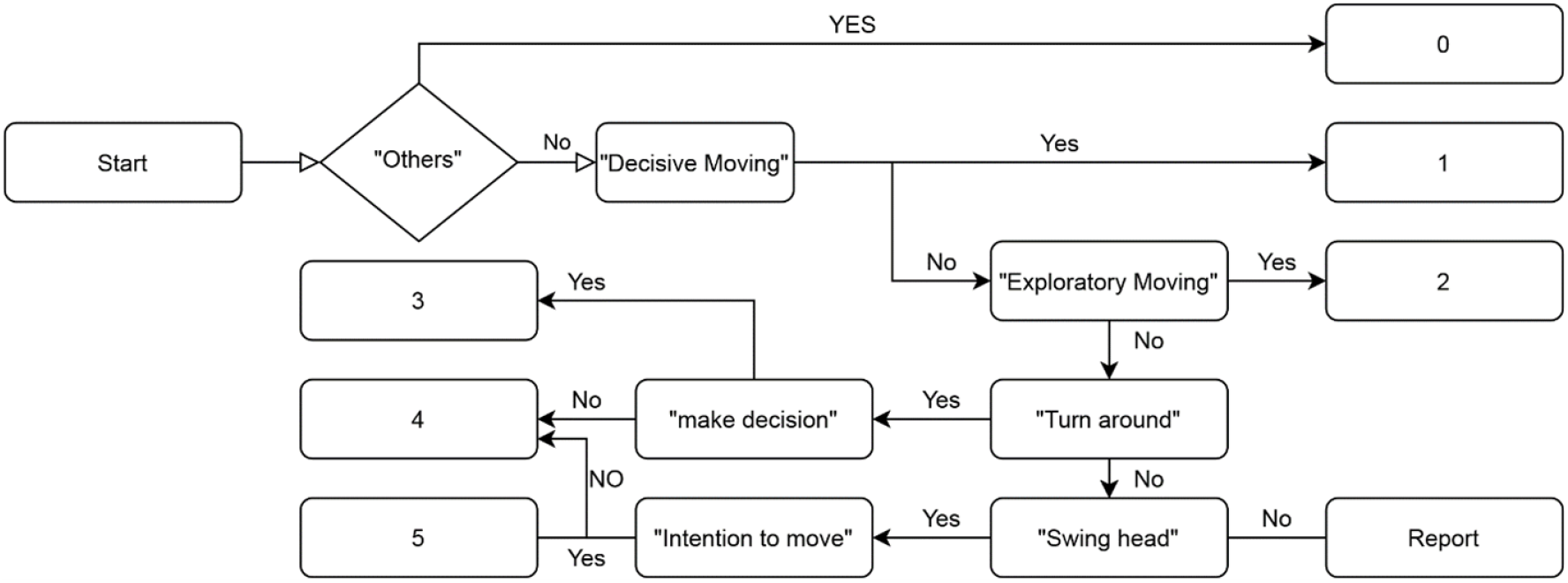
Decision pipeline of uncertainty ratings.

**Table B.2:** Wayfinding tasks.

| Task | Origin | Destination | Description |
| --- | --- | --- | --- |
| Task 1 | Info desk | Elevator | The shortest path included walking through the main entrance hallway with furniture on the left side, then seeing the sign for “Ambulatory Care” with directions to turn right at the first intersection, and then seeing a T-shape intersection with the sign “Medical Imaging” in front. After turning right participants could see the sign of “Main Elevator” at the end of the hallway. |
| Task 2 | Elevator | Nurse station (Unit A) | The shortest path included pressing a button to go up, leaving the elevator and seeing a white wall with the icon “Floor 5,” seeing T-shape intersections on both the right and left, seeing a sign with information about Unit A, and going through the corresponding corridor to reach the destination at a center of an H-shape intersection with a sign “Unit A – Care Station.” |
| Task 3 | Nurse station (Unit A) | Patient room #5A-511 | The shortest path included reading the sign listing patient room numbers, taking the appropriate corridor, and finding the appropriate room in the corridor. |
| Task 4 | Patient room #5A-511 | Elevator | Same environment as described in Tasks 2 and 3. |
| Task 5 | Elevator | Hospital main entrance | Same environment as described in Tasks 2. |
| Task 6 | Hospital main entrance | Ambulatory care reception desk | Same environment as described in Tasks 1 and 2. After seeing the sign “Ambulatory Care” individuals will turn right and see another hallway with the sign “Ambulatory Care Reception Desk” in front of them. |
| Task 7 | Ambulatory care reception desk | Treatment chair #4 (Section C) | The shortest path included seeing three corridors with the large icons “A,” “B,” and “C” on the walls, then going through the appropriate hallway past room number signs on both sides, passing an intersection with information about Clinic C, then reaching the appropriate chair. |
| Task 8 | Treatment chair #4 (Section C) | Back to the ambulatory care reception desk | Same environment as described in Task 7. |
| Task 9 | Ambulatory care reception desk | Cafeteria cashier | The shortest path included seeing the sign in the front corridor describing the direction to Medical Imaging, Cafeteria, and Ambulatory Care, then turning left, reaching a T-intersection, and seeing the cafeteria located to the right. |
| Task 10 | Cafeteria cashier | Hospital main entrance | The shortest path back to the hospital entrance included reaching the main hallway then following a short corridor to the information desk, then turning to the right. |

